# Frizzled-9 activates YAP to rescue simulated microgravity induced osteoblasts dysfunction

**DOI:** 10.1101/2023.03.31.535068

**Authors:** Qiusheng Shi, Jinpeng Gui, Yaxin Song, Jing Na, Jingyi Zhang, Lianwen Sun, Yubo Fan, Lisha Zheng

## Abstract

Long-term space flight will lead to bone loss and osteoblasts dysfunction. The underlying mechanism is still far to reveal. Frizzled-9 (Fzd9) is a Wnt receptor which is essential to osteoblasts differentiation and bone formation. Here we investigate whether Fzd9 plays a role in simulated microgravity (SMG) induced osteoblasts dysfunction. After 1-3 days of SMG, the osteogenic markers were decreased which accompanied the decline of Fzd9 expression. Fzd9 also decreased in the femur of the rats after 3 weeks of hindlimb unloading. Overexpression of Fzd9 will counteract SMG-induced osteoblasts dysfunction. However, Fzd9 overexpression did not affect SMG induced pGSK3β and β-catenin expression or sublocalization. Overexpression of Fzd9 regulates the phosphorylation of Akt and ERK, as well as induces F-actin polymerization to form the actin cap, presses the nuclei, and increases the nuclear pore size, which promotes nuclear translocation of YAP. Our study provides mechanistic insights into the role of Fzd9 triggers actin polymerization and activates mechano-transducer YAP to rescue SMG-mediated osteoblasts dysfunction and indicates Fzd9 as a potential target to restore osteoblast function in bone diseases and space flight.

## Introduction

Mechanical stresses play a crucial role in bone formation and development (Harada & Rodan, 2003; Hsieh & Turner, 2001). Long-term space flight will lead to bone loss (Leblanc *et al*, 1990), bone formation disorder (Morey & Baylink, 1978), bone resorption (Kaplanskii *et al*, 1987), mineralization dysfunction (Zerath *et al*, 1996), and collagen disorganization (Doty *et al*, 1992). Osteoblasts dysfunction is regarded as one of the important reasons for microgravity-induced bone loss. Shi reported that microgravity abrogated primary cilia of osteoblasts and induced inhibition of osteoblastic differentiation and mineralization (Shi *et al*, 2017). Microgravity also decreased the proliferation of osteoblasts (Kumei *et al*, 1996). The expression of osteogenic markers such as collagen I alpha1, alkaline phosphatase (ALP), and osteocalcin was decreased after a 9 days space flight in MG-63 cells (Carmeliet *et al*, 1997). SMG treatment suppresses the expression of PIEZO1 and impairs osteoblast function (Sun *et al*, 2019a). These studies indicated that microgravity would inhibit the cell viability and osteogenesis of osteoblasts.

Increasing researches try to reveal the underlying mechanisms of microgravity-induced osteoblast dysfunction. Rijken demonstrated that microgravity strongly suppressed EGF and PMA-induced c-fos and c-jun expression in A431 cells. PKC signal pathway and actin microfilament organization were sensitive to microgravity (Rijken *et al*, 1994). MAPK pathway was reported to alter the actin dynamics of osteoblast under microgravity (Dai *et al*, 2011). SMG also reduces intracellular-free calcium concentration by inhibiting calcium channels in primary mouse osteoblasts (Sun *et al*, 2019b). PIEZO1-deficient mice are resistant to further bone loss and bone resorption induced by hind limb unloading, demonstrating that PIEZO1 can affect osteoblast-osteoclast crosstalk in response to SMG (Wang *et al*, 2020). However, the mechanism of microgravity regulated osteoblasts is far to reveal.

Fzd9 is a member of the Frizzled family which act as co-receptors with LRP5/6 in Wnt pathway (Wang *et al*, 2016). It was first identified in Williams-Beuren syndrome in 1997 (Schubert, 2009; Wang *et al*, 1997). Fzd9 plays an important role in kidney, brain and bone development (Albers *et al*, 2011; Wang *et al*, 1999). Fzd9-deficient mice displayed a low bone formation due to osteoblasts defect. The mineralization also largely decreased in Fzd9^-/-^ osteoblasts (Albers *et al*., 2011). The new bone formation was impaired in the fracture callus of Fzd9^-/-^ mice (Heilmann *et al*, 2013). However, whether Fzd9 involved in microgravity-induced osteoblast dysfunction was still unknown.

Yes-associated protein (YAP) is an important mechanosensitive co-transcription factor that translocates to the nucleus at higher cellular tension to transduce these mechanical changes in the cell to changes in gene expression (Dupont *et al*, 2011). YAP translocates to the nucleus to control transcriptional regulation of cell fate in response to biophysical cues, such as extracellular matrix (ECM) rigidity (Calvo *et al*, 2013; Chaudhuri *et al*, 2016), strain (Benham-Pyle *et al*, 2015; Elosegui-Artola *et al*, 2016), shear stress (Nakajima *et al*, 2017), SMG (Thompson *et al*, 2020) or adhesive area (Wada *et al*, 2011). This mechanical regulation is accompanied by changes in cytoskeletal F-actin and focal adhesions (Aragona *et al*, 2013; Das *et al*, 2016). One of the most actively investigated signaling pathways that regulate osteoblast functions is the YAP signaling pathway. YAP depleting in osteoblast results in reduced osteogenesis (Brandao *et al*, 2019; Zhu *et al*, 2019). Fzd family is regarded as co-receptors for Wnts in the Wnt/β-catenin signaling pathway which plays a key role in bone remodeling (Lin *et al*, 2009). Recent studies uncovered Wnt ligands promote YAP activation via the alternative Wnt signaling pathway in the process of osteogenic differentiation. Alternative Wnt-YAP signaling consists of Wnt-Fzd/ROR-Gα_12/13_-Rho-LATS1/2-YAP (Park *et al*, 2015). However, the mechanism of Fzd9 regulated YAP activation under SMG has yet to be fully characterized.

In our study, we investigated the role of Fzd9 in regulating osteoblast functions under SMG. The results indicated that Fzd9 expression in osteoblasts was decreased under SMG. Overexpression of Fzd9 could rescue the decrease of osteogenic markers such as ALP, OPN and RUNX2. However, overexpression of Fzd9 had no effect on SMG-induced synthase kinase-3β (GSK3β) phosphorylation and β-catenin expression. Overexpression of Fzd9 rescued osteoblasts dysfunction by regulating YAP activation under SMG. Fzd9 inhibits YAP activity by inhibiting pERK and pAkt to reduce F-actin polymerization and adhesion formation. In addition, Fzd9 inhibited actin cap formation, increased nuclear thickness, and decreased nuclear pore size, which inhibited the nuclear translocation of YAP. Our study indicated that SMG downregulated Fzd9 expression and inhibited YAP activity which induced dysfunction of osteoblasts.

## Results

### Overexpression of Fzd9 rescued SMG-induced osteoblasts dysfunction

Rat osteoblasts were exposed to SMG by a 2D-rotation device for 1-3 days. As shown in Fig 1A-C, we found that the mRNA expression of osteogenic differentiation markers, such as ALP, OPN and RUNX2 was significantly decreased. ALP activity and ALP staining showed the same trend (Fig 1D-F). The above results proved that SMG down-regulated the osteogenic differentiation tendency of osteoblasts. SMG suppressed the gene and protein expression of Fzd9 after exposure to SMG for 3 days (Fig 1G-I). It seemed that Fzd9 had a similar expression mode with osteogenic markers. To investigate the expression pattern of Fzd9 *in vivo*, the rats were hindlimb unloaded for 3 weeks. We detected the bone mineral density of femurs and cortical thickness in rats of the control (Con) and tail suspension (TS) groups. The result showed that the bone mineral density of femurs and cortical thickness of the TS group was significantly less than the Con group after 3 weeks of unloading (Fig 1J-M). We tested the mRNA levels of Fzd9 in the femur and skeleton muscle which highly expressed Fzd9. The results indicated that the Fzd9 gene expression of femur, but not skeleton muscle, declined after hindlimb unloading. It coincided with the in vitro experiment of osteoblasts (Fig 1N). To investigate the role of Fzd9 in SMG-induced osteogenic dysfunction in osteoblasts, we applied a lentivirus system to overexpress Fzd9 in osteoblasts and it was verified by qRT-PCR and western blot (Fig EV1A-C). We discovered that ALP, OPN, and RUNX2 gene expression were markedly increased in the SMG-stimulated osteoblasts after overexpressing Fzd9 (Fig 1O-Q). Additionally, ALP activity and ALP staining in Fzd9 overexpressed osteoblasts were significantly elevated (Fig 1R-T). These results showed that SMG induced osteogenic dysfunction was rescued by Fzd9 in osteoblast.

**Figure 1.**
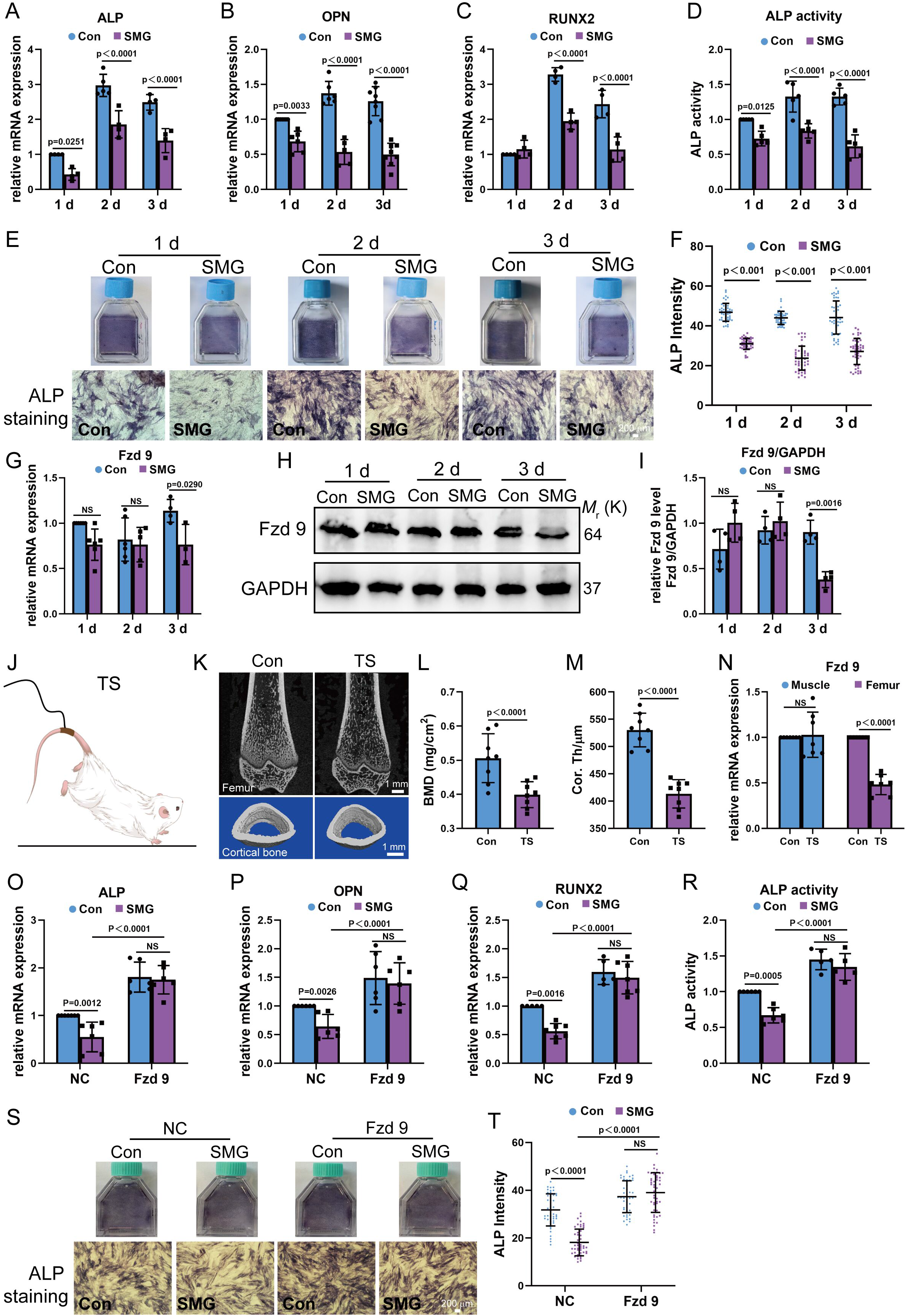
Overexpression of Fzd9 rescued SMG induced osteoblasts dysfunction. A-F The mRNA expression of osteogenesis markers ALP (A, n = 4), OPN (B, n = 6) and RUNX2 (C, n = 4), ALP activity tests (D, n = 5), ALP staining (E, scale bar: 200 mm) and ALP intensity (F, n = 46 for each group) in osteoblasts under 1-3 days of SMG. G-I The mRNA (G, 1 d, n = 6; 2 d, n = 5; 3 d, n = 4) and protein (H and I, n = 4) expression of Fzd9 under 1-3 days of SMG (n ≥ 3). J-N (J, K) 3D micro-CT images of cortical bone of distal femurs isolated from 3 weeks of hindlimb unloading (scale bar: 1mm). Micro-CT analysis of distal femurs from for bone mineral density (L, BMD, n = 8 for each group) and cortical thickness (M, cor. Th, n = 8 for each group) of middle shaft of femurs. (N) The mRNA expression of Fzd9 in skeleton muscle and femur (n = 7 for each group). O-T Osteoblasts transfected with LV5-NC or LV5-Fzd9 were kept under static conditions or subjected to SMG. ALP (O, n = 6), OPN (P, n = 6), RUNX2 (Q, n = 6) gene expression, ALP activity tests (R, n = 5), ALP staining (S, scale bar: 200 μm) and ALP intensity (T, n = 47 for each group) were presented. Data information: All graphs show mean ± s.d. n reflects the number of biological replicates, which are summarized from at least three independent experiments. p values were obtained using two-way ANOVA followed by Tukey’s post hoc test (A, B, C, D, F, N, O, P, Q, R, T) or two-tailed Student’s t tests (L, M).

### Fzd9 rescued SMG-induced osteoblasts dysfunction did not mediated by Wnt/β-catenin pathway

Fzd family is regarded as co-receptors for Wnts in the Wnt/β-catenin signaling pathway. Whether Fzd9 regulated osteoblasts by Wnt/β-catenin signaling pathway under SMG is still unknown. We tested two key factors in the canonical Wnt pathway, GSK3β and β-catenin in osteoblasts after SMG. As shown in Fig 2A and B, the nuclear localization of β-catenin was significantly inhibited during the 3 days of SMG. Meanwhile, SMG increased the GSK3β phosphorylation and decreased β-catenin expression (Fig 2C-E). It demonstrated that SMG inhibited Wnt/β-catenin signaling. Overexpressed of Fzd9 did not affect β-catenin nuclear localization (Fig 2F and G), nor the pGSK3β and β-catenin induced by SMG (Fig 2H-K). These results indicated that Fzd9 regulated osteogenesis did not mediate by Wnt/β-catenin pathway under SMG.

**Figure 2.**
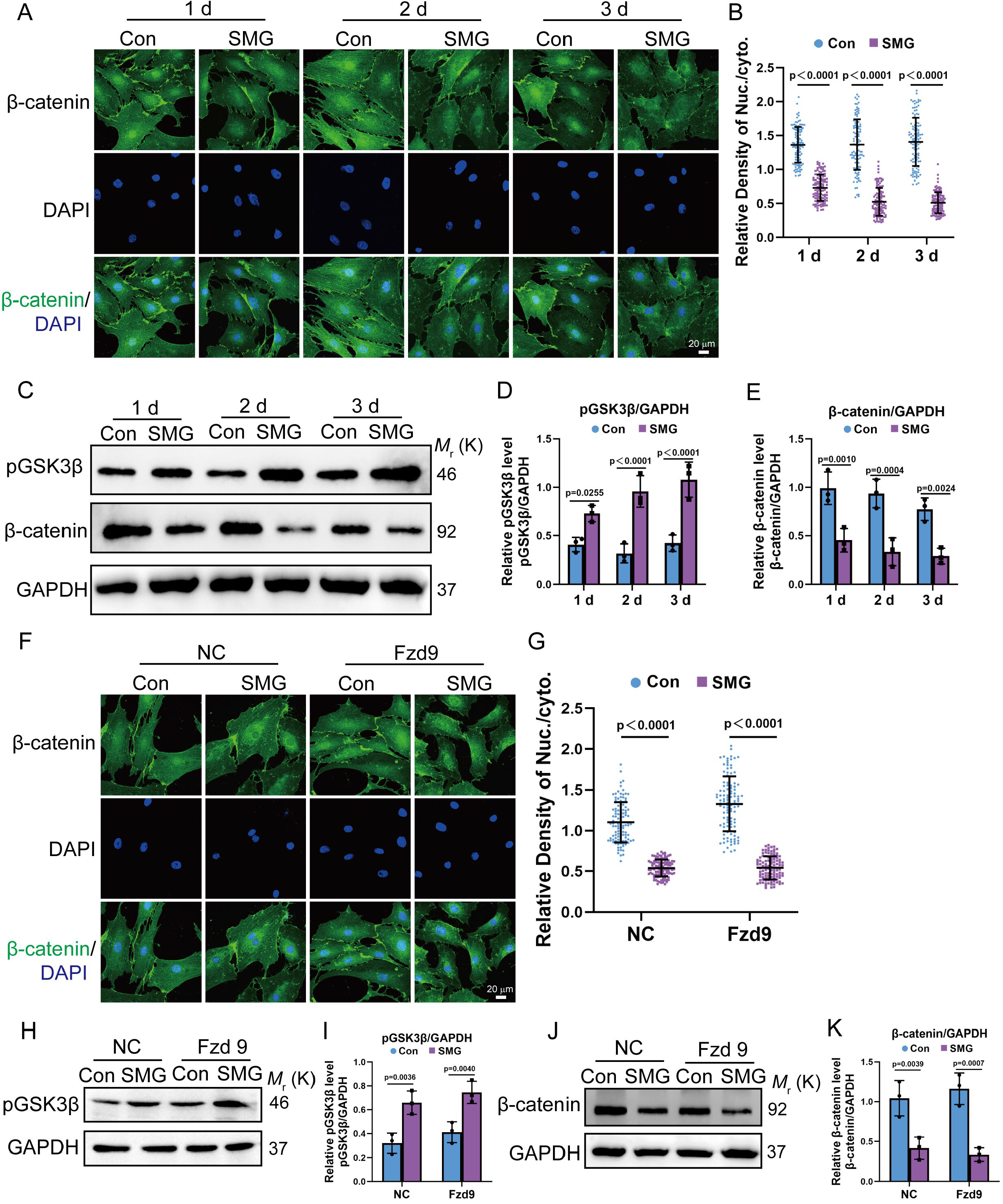
Fzd9 rescued SMG-induced osteoblasts dysfunction did not mediated by Wnt/β-catenin pathway. A, B (A) Osteoblasts under static conditions or after SMG for 1, 2 or 3 days were fixed and immunostained with anti-β-catenin (green) antibody together with DAPI (blue). Scale bars = 20 μm. (B) Quantification of nuclear relative to cytoplasmic fluorescent intensity of β-catenin in static or SMG (n = 110 for each group). C-E (C) pGSK3β and β-catenin protein expressions on static or SMG in osteoblasts were detected by Western blotting. pGSK3β/GAPDH (D, n = 3) and β-catenin/GAPDH (E, n = 3) intensity ratio were shown. F-K Osteoblasts transfected with LV5-NC or LV5-Fzd9 were kept under static conditions or subjected to SMG and β-catenin (green) localization (F, Scale bars = 20 μm), quantification of nuclear relative to cytoplasmic fluorescent intensity (G, n = 109 for each group), pGSK3β/GAPDH expression (H, I, n = 3) and β-catenin/GAPDH expression (J, K, n = 3) were presented. Data information: All graphs show mean ± s.d. n reflects the number of biological replicates, which are summarized from at least three independent experiments. All p values were obtained using two-way ANOVA followed by Tukey’s post hoc test.

### Fzd9 regulated SMG-induced osteoblasts dysfunction by regulating YAP

YAP is a mechanotransducer and plays an important role in cell proliferation, migration, and osteogenic differentiation. We investigated their role in SMG-induced osteoblasts dysfunction. As shown in Fig 3A and B, the nuclear localization of YAP was significantly inhibited during 1-3 days of SMG. The YAP target genes such as CTGF and CYR61 were also inhibited by SMG (Fig 3C and D). These data indicated that SMG inhibited YAP activity. When we knocked down YAP (Fig EV2A and B), the osteogenesis markers were further decreased no matter in the static group or SMG group. And there was no obvious difference between static control with SMG (Fig 3E-G). In addition, the ALP activity and ALP staining results confirmed it (Fig 3H-J). It indicated that YAP was vital to osteogenesis in osteoblasts. When Fzd9 was overexpressed, SMG-induced YAP nuclear expelling, as well as CTGF and CRY61 downregulation, were enhanced (Fig 3K-N). These results demonstrated that YAP involved in SMG-induced osteoblasts dysfunction. Overexpression of Fzd9 could elevate YAP activity to rescue SMG-induced osteogenic depression.

**Figure 3.**
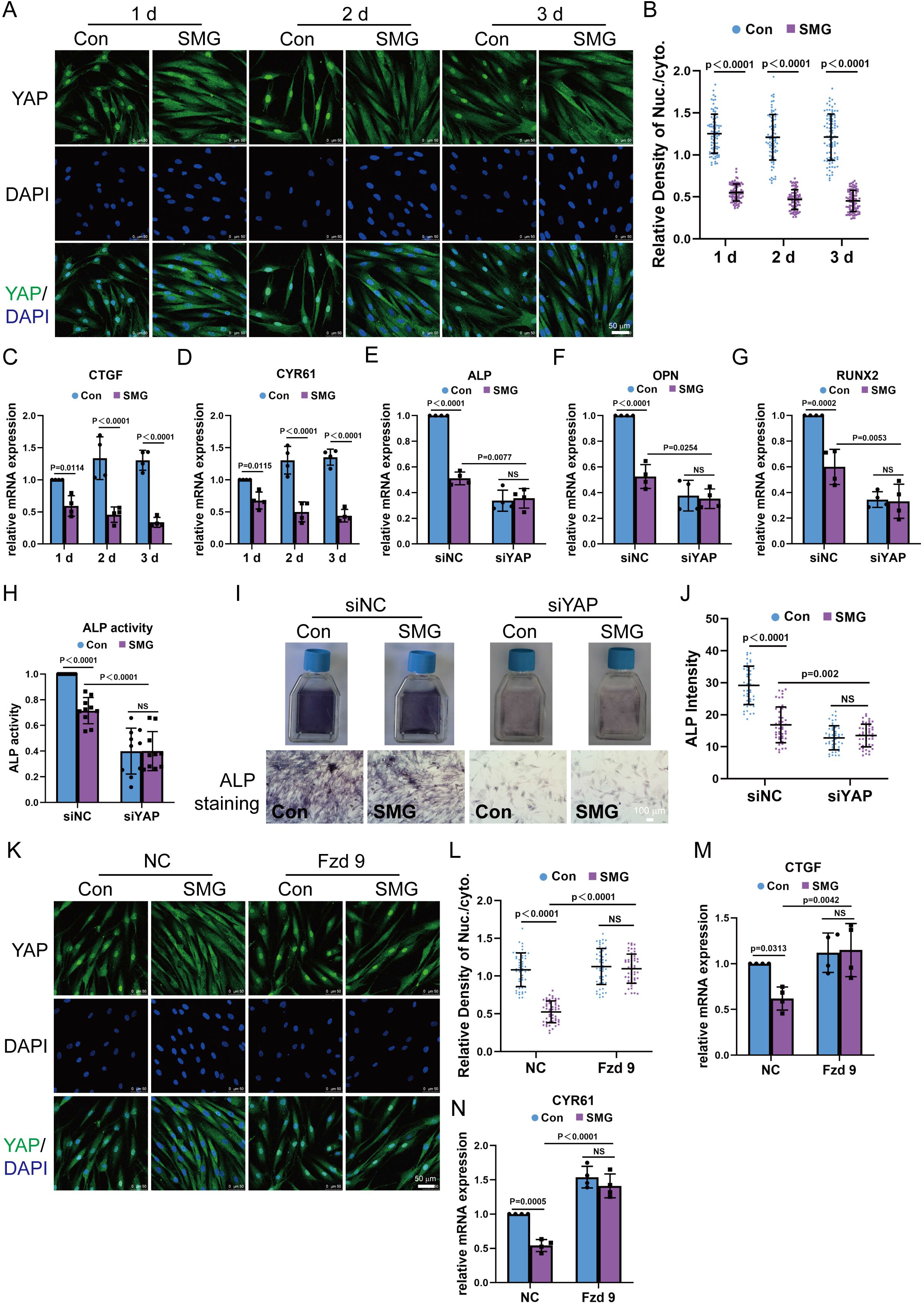
Fzd9 regulated SMG-induced osteoblasts dysfunction by regulating YAP. A-D (A) Osteoblasts under static conditions or after SMG for 1, 2 or 3 days were fixed and immunostained with anti-YAP (green) antibody together with DAPI (blue). Scale bars = 50 μm. (B) Quantification of nuclear relative to cytoplasmic fluorescent intensity of YAP in static or SMG (n = 85 for each group). qPCR of CTGF (C, n = 4) and CYR61(D, n = 4) gene expression under static conditions or SMG. E-J Osteoblasts transfected with control siRNA or YAP siRNAs and subjected to SMG. ALP (E, n = 4), OPN (F, n = 4), RUNX2 (G, n = 4) gene expression, ALP activity tests (H, n = 10), ALP staining (I, scale bar: 200 μm) and ALP intensity (J, n = 47 for each group) were presented. K-N Osteoblasts transfected with LV5-NC or LV5-Fzd9 were kept under static conditions or subjected to SMG and YAP (green) localization (K, scale bar: 50 μm), quantification of nuclear relative to cytoplasmic fluorescent intensity (L, n = 52 for each group), CTGF (M, n = 4) and CYR61 (N, n = 4) gene expression were presented. Data information: All graphs show mean ± s.d. n reflects the number of biological replicates, which are summarized from at least three independent experiments. All p values were obtained using two-way ANOVA followed by Tukey’s post hoc test.

### Fzd9 regulated SMG-induced decrease of F-actin polymerization and focal adhesion formation

Remarkably, the multiple regulatory inputs that determine YAP activity converge on the actin cytoskeleton and focal adhesion. To clarify cytoskeletal organization, osteoblasts were grown under normal gravity and SMG conditions were stained with F-actin and G-actin at days 1, 2 and 3. The cells grown under SMG conditions presented less F-actin and more G-actin (Fig 4A-C). Moreover, paxillin, a focal adhesion protein, was also analyzed at these time points in cells under SMG. The immunofluorescence results showed that SMG decreased the intensity and length of paxillin (Fig 4D-F). These data indicate that SMG could decrease F-actin polymerization and focal adhesion formation. To investigate the role of cytoskeleton in SMG-induced YAP localization, cytochalasin D (Cyto D) was used to disturb actin filament polymerization. The results indicated that the impaired actin filaments accorded with nuclear expulsion of YAP under SMG. On the contrary, when cells were incubated with jasplakinolide (Jasp), a reagent which promotes actin polymerization, the SMG-induced nuclear localization of YAP increased (Fig EV3A and B), indicating that the integrity of actin filaments played an important role in SMG-induced YAP localization. To investigate the role of Fzd9 in SMG decreased F-actin polymerization and focal adhesion formation, we overexpressed Fzd9 in osteoblasts and exposed osteoblasts to SMG for 3 days. We found that F-actin intensity decreased after 3 days of SMG companying with upregulated G-actin intensity was rescued by Fzd9 overexpression (Fig 4G-I). Paxillin intensity and length were enhanced when Fzd9 was overexpressed (Fig 4J-L). These results demonstrated that Fzd9 involved in SMG decrease F-actin polymerization and focal adhesion formation.

**Figure 4.**
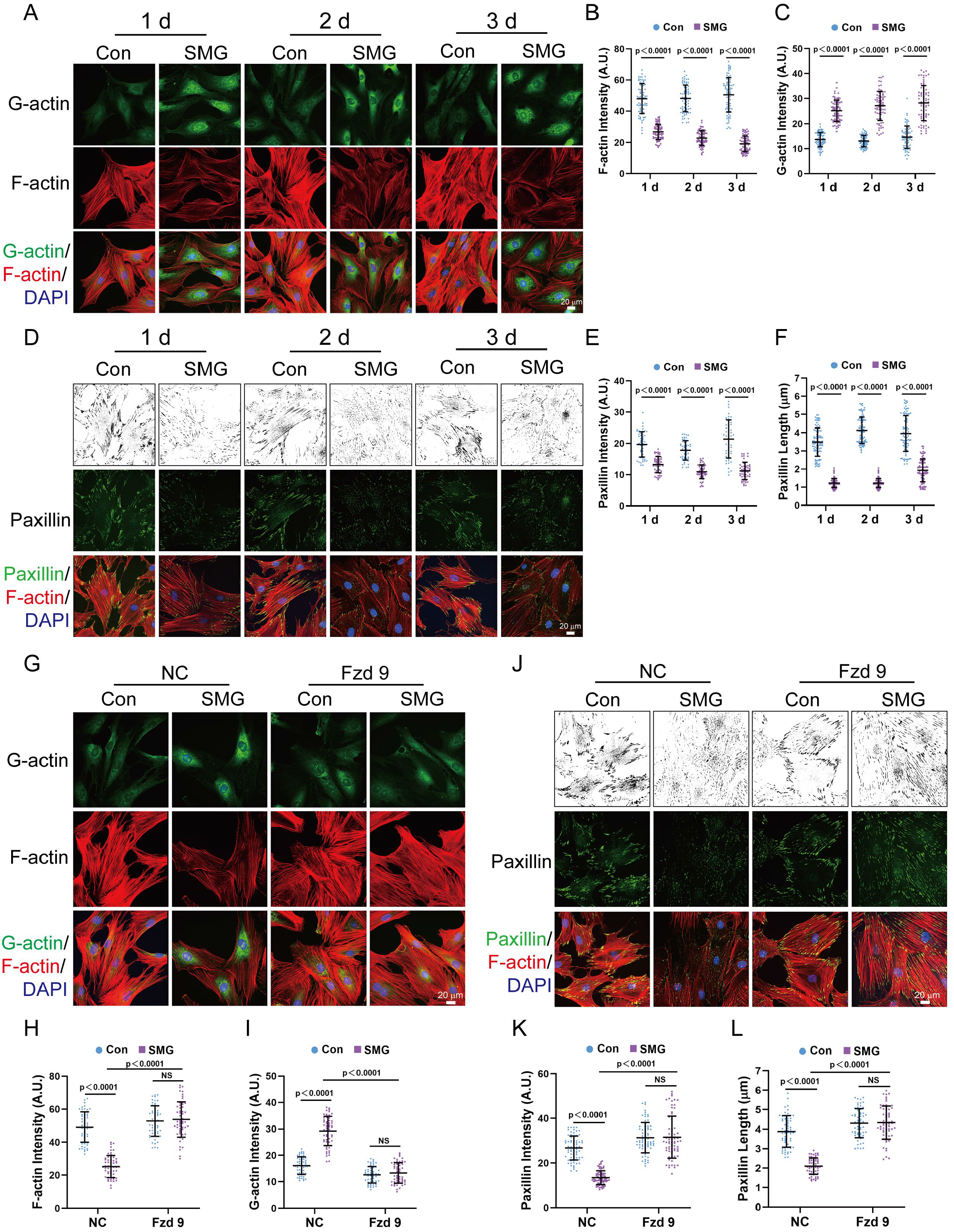
Fzd9 regulated SMG-induced decrease of F-actin polymerization and focal adhesion formation. A-C (A) Osteoblasts under static conditions or after SMG for 1, 2 or 3 days were fixed and immunostained with rhodamine phalloidin (F-actin, red), DNase I 488 (G-actin, green) together with DAPI (blue). Scale bars = 20 μm. Quantification of cell density of F-actin (B, n = 74 for each group) and cell density of G-actin (C, n = 74 for each group) in static or SMG osteoblasts. D-F (D) Osteoblasts under static conditions or after SMG for 1 day, 2 days or 3 days were fixed and immunostained with rhodamine phalloidin (F-actin, red), anti-paxillin (green) antibody together with DAPI (blue). Scale bars = 20 μm. Quantification of paxillin intensity (E, n = 52 for each group) and paxillin length (F, n = 91 for each group) in static or SMG osteoblasts. G-I (G) Osteoblasts transfected with LV5-NC or LV5-Fzd9 were kept under static conditions or subjected to SMG and immunostained with rhodamine phalloidin (F-actin, red), DNase I 488 (G-actin, green) together with DAPI (blue). Scale bars = 20 μm. Quantification of cell density of F-actin (H, n = 55 for each group) and cell density of G-actin (I, n = 55 for each group) in static or SMG osteoblasts. J-L (J) Osteoblasts transfected with LV5-NC or LV5-Fzd9 were kept under static conditions or subjected to SMG and immunostained with rhodamine phalloidin (F-actin, red), anti-paxillin (green) antibody together with DAPI (blue). Scale bars = 20 μm. Quantification of paxillin intensity (K, n = 72 for each group) and paxillin length (L, n = 61 for each group) in static or SMG osteoblasts. Data information: All graphs show mean ± s.d. n reflects the number of biological replicates, which are summarized from at least three independent experiments. All p values were obtained using two-way ANOVA followed by Tukey’s post hoc test.

### Fzd9 affected ERK/Akt to regulate SMG decreased F-actin polymerization and focal adhesion formation

It has been reported that the presence of Fzd9 in osteoblasts is required to induce noncanonical Wnt signaling pathways, and they are in full agreement with the positive influence of increased ERK and Akt signaling on bone formation (Albers *et al*., 2011). Meanwhile, ERK and Akt are key factors that regulate actin polymerization (Hollosi *et al*, 2022; Xue & Hemmings, 2013). Interestingly, our results indicated that pERK decreased by 1-3 days of SMG (Fig 5A and B). Furthermore, the levels of pAkt also were downregulated by 1-3 days of SMG (Fig 5C and D). SMG-inhibited pAkt expression was further decreased when we used the ERK inhibitor PD (Fig 5E and F). When osteoblasts were preincubated with the ERK inhibitor or Akt inhibitor and loaded with 3 days of SMG, polymerization of F-actin decreased compared with the control (Fig 5G-L). Besides, the paxillin intensity and length were significantly restrained (Fig 5M-R). Thereby ERK/Akt regulate the effects of SMG on F-actin polymerization and focal adhesion formation. To demonstrate the relationship between Fzd9 and ERK/Akt, we overexpressed Fzd9 in osteoblasts and exposed osteoblasts to SMG for 3 days. The results indicated that SMG decreased pERK, pAkt levels were rescued by Fzd9 (Fig 5S-V). These results demonstrated that Fzd9 involved in ERK/Akt to regulate SMG decreased F-actin polymerization and focal adhesion formation in osteoblasts.

**Figure 5.**
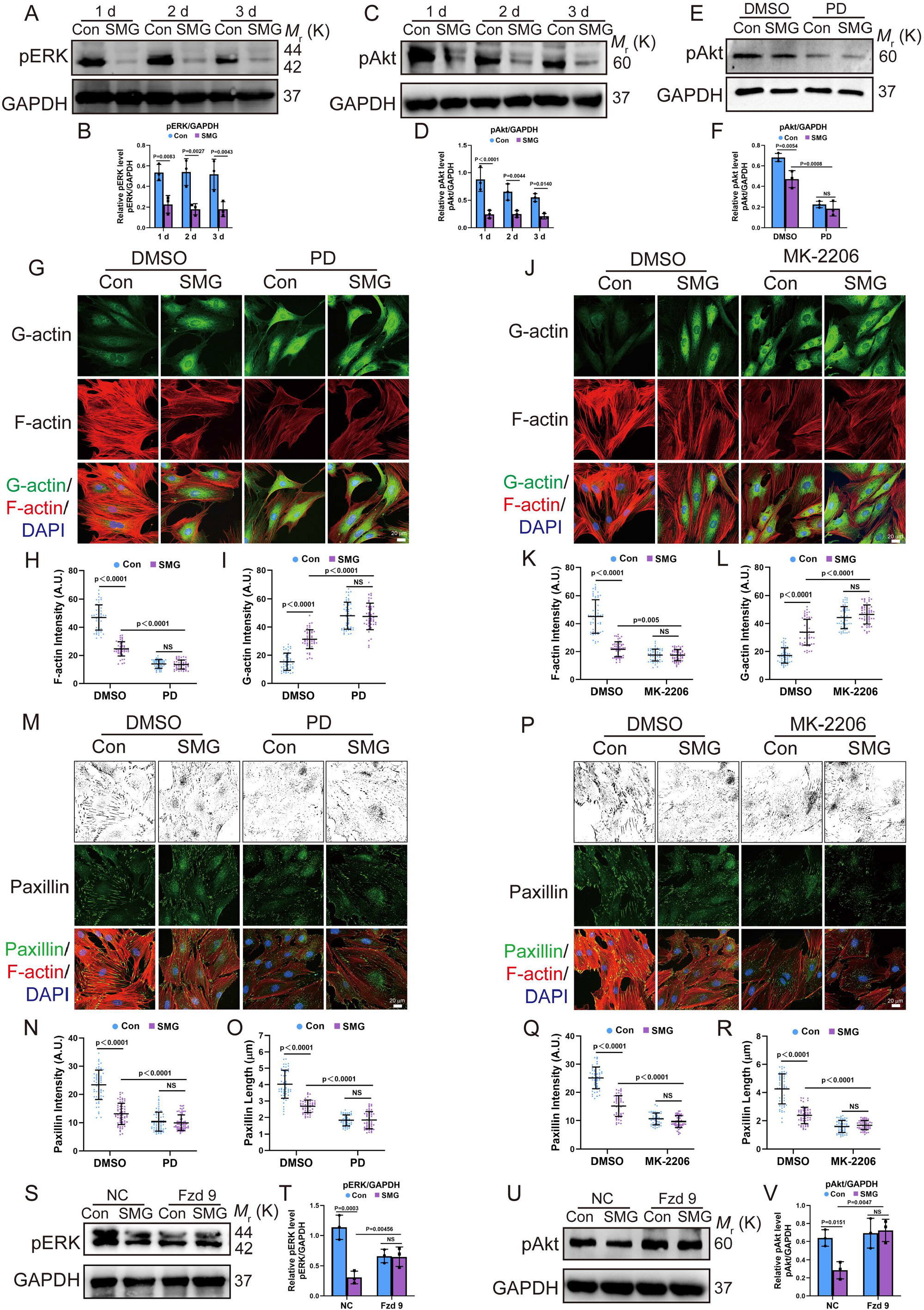
Fzd9 affected ERK/Akt to regulate SMG decreased F-actin polymerization and focal adhesion formation. A-D pERK (A) and pAkt (C) protein expressions on static or SMG in osteoblasts were detected by Western blotting. pERK/GAPDH (B, n = 3) and pAkt/GAPDH (D, n = 3) intensity ratio were shown. E, F Osteoblasts preincubated with DMSO or ERK pathway inhibitor PD (10 μm, 2 h) and under static conditions or subjected to SMG for 3 days. pAkt protein expressions (E) were detected by Western blotting. pAkt /GAPDH intensity ratio were shown (F, n = 3). G-L (G, J) Osteoblasts preincubated with DMSO, ERK pathway inhibitor PD98059 (PD, 10 μm, 2 h) or Akt pathway inhibitor MK-2206 (2.5 μm, 24 h) and under static conditions or subjected to SMG for 3 days. Osteoblasts were fixed and immunostained with rhodamine phalloidin (F-actin, red), DNase I 488 (G-actin, green) together with DAPI (blue). Scale bars = 20 μm. Quantification of cell density of F-actin (H, n = 51 for each group, K, n = 55 for each group) and cell density of G-actin (I, n = 51 for each group, L, n = 55 for each group) in static or SMG osteoblasts. M-R (M, P) Osteoblasts preincubated with DMSO, ERK pathway inhibitor PD98059 (PD,10 μm, 2 h) or Akt pathway inhibitor MK-2206 (2.5 μm, 24 h) and under static conditions or subjected to SMG for 3 days. Osteoblasts were fixed and immunostained with rhodamine phalloidin (F-actin, red), anti-paxillin (green) antibody together with DAPI (blue). Scale bars = 20 μm. Quantification of paxillin intensity (N, n = 64 for each group, Q, n = 58 for each group) and paxillin length (O, n = 51 for each group, R, n = 51 for each group) in static or SMG osteoblasts. S-V Osteoblasts transfected with LV5-NC or LV5-Fzd9 were kept under static conditions or subjected to SMG for 3days. pERK (S) and pAkt (U) protein expressions were detected by Western blotting. pERK/GAPDH (T, n = 3) and pAkt/GAPDH (V, n = 3) intensity ratio were shown. Data information: All graphs show mean ± s.d. n reflects the number of biological replicates, which are summarized from at least three independent experiments. All p values were obtained using two-way ANOVA followed by Tukey’s post hoc test.

### Fzd9 regulated SMG-induced nuclear morphology

Recently, nuclear flattening was proposed to generally increase size of nuclear pores and has been reported to regulate YAP activity. In our study, when SMG was loaded on osteoblasts for 1-3 days, the actin cap declined, the nuclear thickness increased, the nuclear area and nuclear volume depressed within 1-3 days (Fig 6A-E). The total fluorescence intensity of actin cap and the nuclear thickness showed a high correlation, demonstrating that SMG inhibited actin cap expansion and raised nuclei (Fig 6F). The N/C of the YAP ratio was highly correlated with the nuclear thickness, which indicated that the nuclear thickness affected YAP nuclear localization (Fig 6G). We further investigated the effect within 1-3 days of SMG on the nuclear pore. The TEM results indicated that 1-3 days of SMG decreased the nuclear pore size (Fig 6H and I). The nuclear export inhibitor (LMB) blocked 3 days of SMG-induced YAP export, which showed that nuclear pore export was essential for SMG-induced YAP transport (Fig EV4A and B). The Fzd9 overexpressed osteoblasts were exposed to SMG for 3 days. We found that actin cap in Fzd9-overexpressed osteoblasts was significantly elevated. Overexpression of Fzd9 counteract SMG-induced nuclear thickness increased which did not response to SMG regulation (Fig 6J-N). When we overexpression Fzd9, SMG-induced nuclear pore size decreased was blocked (Fig 6O-P). These results demonstrated that Fzd9 regulated SMG inhibited actin cap formation, increased nuclear thickness, and decreased nuclear pore size, which decreased nuclear translocation of YAP.

**Figure 6.**
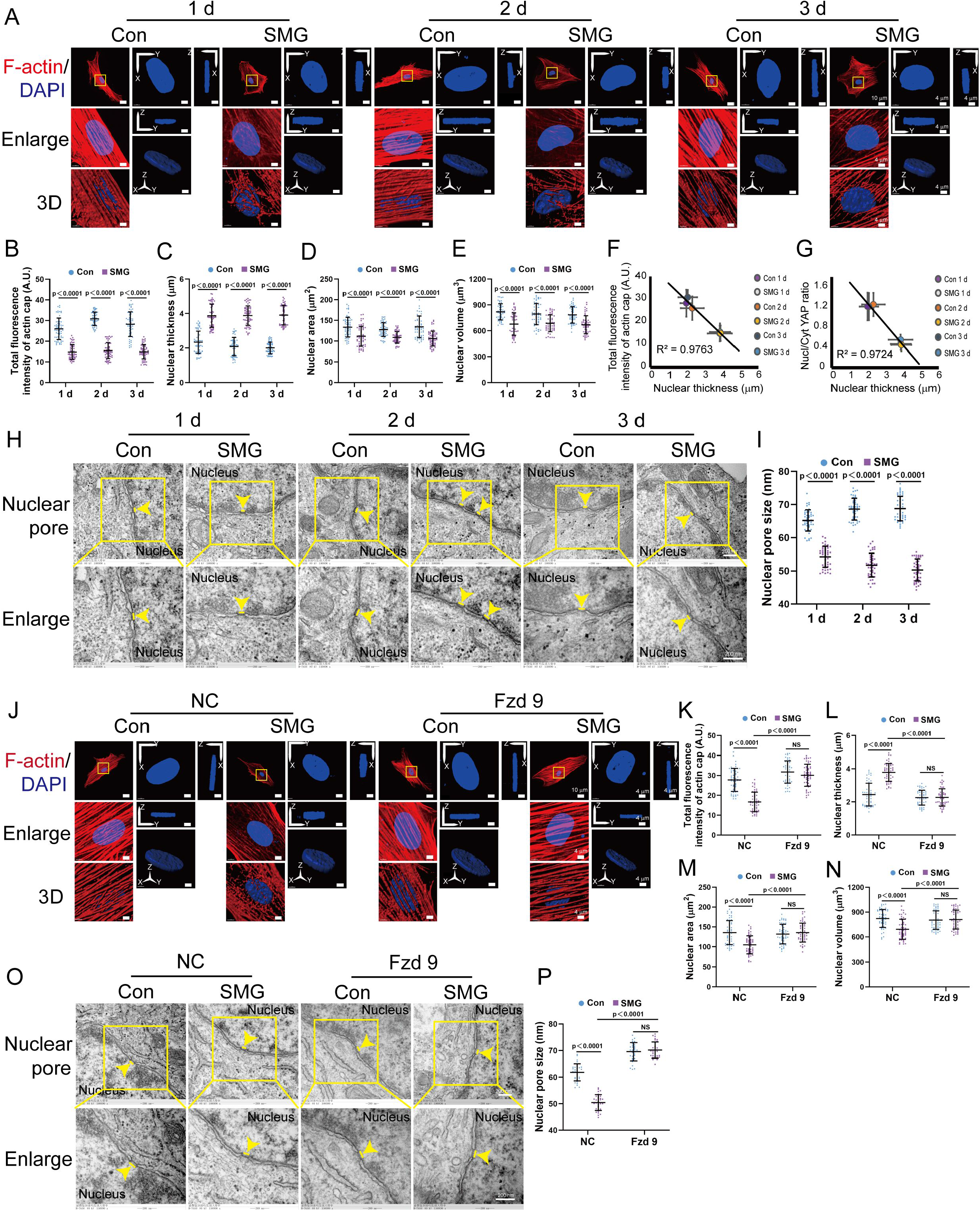
Fzd9 regulated SMG-induced nuclear morphology. A-E (A) Osteoblasts under static conditions or after SMG for 1, 2 or 3 days were fixed and immunostained with rhodamine phalloidin (F-actin, red) together with DAPI (blue). Scale bars = 10 μm. F-actin organization in the apical region of the nucleus were given. 3D rendering illustrate indicated the presence of the perinuclear actin cap. Scale bars = 4 μm. Nuclear morphology of DAPI-stained nuclei (blue) indicates 3D nuclear shape, where maximum intensity projection onto the XY-plane was performed using upper hemispheres of the 3D reconstructed nuclei. The cross-sectional side view was captured along the XZ-plane and YZ-plane crossing the center of the nucleus. Scale bars = 4 μm. Yellow boxes indicate the regions that were shown as a magnified view. Total fluorescence intensity of actin cap (B, n = 58 for each group), Nuclear thickness (C, n = 58 for each group), the projected nuclear area onto the XY-plane (D, n = 58 for each group) and the volume of the 3D reconstructed nuclei (E, n = 58 for each group) were shown. F, G (F) Total fluorescence intensity of actin cap versus nuclear thickness for the conditions in (B) and (C). The dashed line shows a linear fit to the data (R^2^, squared correlation coefficient). (G) Nuc/Cyt YAP ratio versus nuclear thickness for the conditions in Fig. 3B and (C). The dashed line shows a linear fit to the data (R^2^, squared correlation coefficient). H, I (H)TEM images of nuclear pores of osteoblasts under static conditions or after SMG for 1 day, 2 days or 3 days. Yellow arrows indicated nuclear pores. Scale bars = 200 nm. (I) Quantification of Nuclear pore size (n = 45 roi per condition nuclear pores from ≥ 15 cells per condition) in static or SMG osteoblasts. J-N (J) Osteoblasts transfected with LV5-NC or LV5-Fzd9 were kept under static conditions or subjected to SMG and immunostained with rhodamine phalloidin (F-actin, red) together with DAPI (blue). Yellow boxes indicate the regions that were shown as a magnified view. Total fluorescence intensity of actin cap (K, n = 53 for each group), Nuclear thickness (L, n = 53 for each group), the projected nuclear area onto the XY-plane (M, n = 53 for each group) and the volume of the 3D reconstructed nuclei (N, n = 53 for each group) were shown. O, P (O) TEM images of nuclear pores of osteoblasts transfected with LV5-NC or LV5-Fzd9 were kept under static conditions or subjected to SMG. Yellow arrows indicated nuclear pores. (P) Quantification of Nuclear pore size (n = 31 roi per condition nuclear pores from ≥ 15 cells per condition) in static or SMG osteoblasts. Data information: All graphs show mean ± s.d. n reflects the number of biological replicates, which are summarized from at least three independent experiments. All p values were obtained using two-way ANOVA followed by Tukey’s post hoc test.

## Discussion

Microgravity during spaceflight will result in deleterious bone architectural changes with a decrease in bone volume and density, and further lead to bone loss. The microgravity caused bone loss and osteoblasts dysfunction is an inevitable outcome for astronauts in the long duration of spaceflight. However, the underlying mechanism has not been well elucidated. Our current study found that Fzd9 acts in osteoblasts as a regulator of bone remodeling via sensing SMG. Overexpression Fzd9 could rescue SMG-induced osteogenic dysfunction in osteoblasts. Our mechanism study demonstrates that Fzd9 regulates YAP signaling, which in turn regulates the expression of osteogenic markers in osteoblasts under SMG.

The expression of Fzd9 showed the correlation with the depression of osteogenic markers, such as ALP, OPN, and RUNX2 in osteoblasts under SMG. It was reported that Fzd9 promoted osteoblast differentiation and new bone formation in fracture healing. Knocking out Fzd9 did not affect osteoclasts (Albers *et al*., 2011; Heilmann *et al*., 2013). Fzd9 expression increased during early osteoblast differentiation in 0-5 days (Albers *et al*., 2011). Our results showed that the expression of Fzd9 was depressed by 3 days of SMG. The Fzd9 expression of femur in hindlimb unloading rat was depressed too. It indicated that Fzd9 was sensitive to mechanical loading. We further investigated the potential role of Fzd9 in SMG-induced bone loss. Overexpression of Fzd9 could increase SMG-induced osteogenic depression in osteoblasts. Our data provided evidence that Fzd9 could regulate osteogenic function under SMG in osteoblasts.

Wnt proteins are morphogens that elicit diverse receptor-mediated signaling pathways to control development and tissue homeostasis. Canonical Wnt signaling acts through β-catenin/TCF transcriptional activity (referred to as ‘‘Wnt/β-catenin signaling’’). Noncanonical Wnt signaling mediates biological responses that do not involve β-catenin/TCF activity (referred to as ‘‘alternative Wnt signaling’’) (Logan & Nusse, 2004). Fzd9 is a member of the Frizzled family which is receptors with LRP5/6 in Wnt pathway. Our results indicated that the canonical Wnt/β-catenin pathway was inhibited by SMG. which consisted of previous studies (Norsk, 2005). Overexpress of Fzd9 did not affect SMG-induced pGSK3β and β-catenin expression or sublocalization. It indicated that SMG-induced Fzd9 decrease was not associated with Wnt/β-catenin pathway. It echoed the previous study that canonical Wnt signaling was not impaired in the absence of Fzd9 in osteoblasts (Albers *et al*., 2011).

YAP is a mechanosensitive transcriptional regulator with a major role in differentiation (Zhang *et al*, 2022). To investigate the underlying mechanism of Fzd9 under SMG, we further studied YAP which was associated with osteogenesis. In this study, we investigated the mechanism underlying SMG induced osteoblasts dysfunction. We observed that the nuclear localization of YAP was significantly inhibited during 1-3 days of SMG. Additionally, we found that YAP target genes such as CYR61 and CTGF were also inhibited by SMG. Knocking down YAP suppressed the tendency of osteogenic differentiation in osteoblasts. These findings demonstrate that YAP functions as an important regulator to promote osteogenic differentiation in SMG inhibited osteogenic differentiation. These results are similar with previous reports that SMG inhibited osteogenic differentiation of mesenchymal stem cells (MSCs) via depolymerizing F-actin to impede TAZ nuclear translocation (Chen *et al*, 2016). Overexpression of Fzd9 could elevate YAP activity to rescue SMG-induced osteogenic depression. Consistent with our results, Fzd10 shows homology to Fzd9 and was mediates G alpha (12/13) activation-dependent induction of YAP transcriptional activity (Hot *et al*, 2017).

Increasing studies have demonstrated that YAP activity is regulated by the conformation and tension of the F-actin cytoskeleton, which in turn depends primarily on the substrate to which cells adhere (Das *et al*., 2016). Our results indicate that disrupting cytoskeleton integrity prevented SMG-mediated YAP nuclear localization. However, promoting F-actin polymerization increased SMG-mediated YAP nuclear transport. Our study found that SMG could decrease F-actin polymerization and focal adhesion formation, which may explain why YAP was transported to the cytoplasm. Our findings agreement with previous studies reporting that human BMSCs in SMG showed F-actin depolarization (Montagna *et al*, 2022). ERK/Akt pathway is a critical mediator of actin reorganization in different types of cells (Menu *et al*, 2004; Xue & Hemmings, 2013). We establish that the ERK and Akt signaling pathways are important links between F-actin cytoskeleton remodeling and YAP activation. SMG could decrease pERK and pAkt. The inhibition of ERK decreased SMG-induced phosphorylation of Akt, as well as F-actin density, indicating that ERK is the upstream regulator of F-actin polymerization. Albers et al., showed that the Wnt5a-induced phosphorylation of ERK and Akt was less pronounced in Fzd9^-/-^ cells, thereby providing evidence for impaired noncanonical Wnt signaling in the absence of Fzd9 (Albers *et al*., 2011). Similarly, in our study, we found that SMG-induced inhibition of pERK and pAkt, as well as actin filament disorganization, was reversed by overexpression of Fzd9. Thereby, it showed the function of Fzd9 to rescue F-actin polymerization to regulate YAP activity.

The perinuclear actin cap that physically connects the apical side of the nucleus to the basal side of the cell (via actin cap associated focal adhesions), where the actin cap serves as a conduction path of the mechanical signal and simultaneously discharges the stresses transferring onto the nucleus (Kim *et al*, 2013; Kim *et al*, 2017). It has been reported that actin cap induced nuclear flattening can increase nuclear pore permeability and promote YAP nuclear import (Elosegui-Artola *et al*, 2017; Shi *et al*, 2022). In our study, we found that the actin cap decreased by SMG. Furthermore, the nuclear thickness was correlated with the actin cap formation, indicating that SMG inhibited actin cap formation, increased nuclear thickness, and decreased nuclear pore size, which inhibited nuclear translocation of YAP. Consistent with our model, Khatau et al., showed that when the actin cap of adherent cells is disrupted with a low-dose treatment of the F-actin depolymerizing drug latrunculin B or by disruption of the LINC complexes, nuclear lobulation ensues and the nucleus can bulge to almost twice its original height (Khatau *et al*, 2009). In addition, Fzd9 exerted its regulatory effects through rescue actin cap formation, nuclear thickness and nuclear pore size. Here, we demonstrate the link between Fzd9 and YAP under SMG and further show that Fzd9 could also reduce nuclear thickness-induced YAP nuclear translocation by rescuing the actin cap. Our findings reveal a novel mechanosensing mechanism Fzd9-induced YAP activation.

In conclusion, our data suggest that SMG reduced the expression of Fzd9 accompanying the dysfunction of osteoblasts. Overexpression of Fzd9 could rescue SMG-induced osteogenic impairment through YAP. Fzd9 inhibited the phosphorylation of ERK and Akt, as well as decreased F-actin polymerization and focal adhesion formation to inhibite YAP activity. In addition, Fzd9 regulates SMG inhibited actin cap formation, increased nuclear thickness, and decreased nuclear pore size, which inhibited nuclear translocation of YAP. Our study indicated Fzd9 as a positive regulator in SMG-induced osteogenic dysfunction in osteoblasts. It contributes to a better understanding of SMG regulated osteogenesis and provides a potential target to restore osteoblast function in bone diseases and space flight.

## Materials and Methods

### Primary Rat Osteoblasts Isolation and Culture

Primary rat osteoblasts were isolated from the calvarias of newborn Sprague-Dawley rats. Rat calvarias were dissected from adherent tissue and washed twice with PBS. The calvarias were shredded to 1 mm × 1 mm pieces and digested for 30 min at 37 °C in Dulbecco’s modifiation of Eagle’s medium (DMEM, Gibico, USA) with 3 mg/ml collagenase I (Sigma-Aldrich, Germany) and 0.25% trypsin (Sigma-Aldrich) mixture. All procedures were approved by the Biological and Medical Ethics Committee of Beihang University. The cells were centrifuged, re-suspended in DMEM with 10% FBS. The cells were cultured in a humidified atmosphere of 5% CO_2_ in air at 37 °C. The medium was changed every 3 days. Upon 80–90% confluency, cells were passaged using 0.25% trypsin, and immuno-localization ALP revealed that all cells were stained positive. All osteoblasts used in this study were 3 to 6 passages.

### Hindlimb Unloading Rat Model

Hindlimb unloading model was used as a bone loss model induced by weightlessness which had been described previously (Sun *et al*, 2011). All the experimental procedures were approved by Biological and Medical Ethics Committee of Beihang University. Briefly, Female Sprague-Dawley (SD) rats, 8 weeks age and body weight ranged 170–190 g, were used for this experiment. After adapting to the laboratory conditions for several days, SD rats were randomly divided into two groups: tail suspension (TS) and control (Con). Rats in TS were individually caged and suspended by the tail at a ∼30° angle to the floor with only the forelimbs touching the floor. Rats in control group were under the same condition without suspending. After 3 weeks of tail suspension, rats were anesthetized for *in vivo* scanning by micro-CT and then sacrificed. Femurs were processed for micro-CT examination. The following parameters were used: scanning voltage, 70 kV; current, 200 μA; exposure time, 300 ms; and thickness, 15 μm. Meanwhile, bone mineral density (BMD) of femurs and cortical thickness (cor, Th), were measured using built-in software after 3D reconstruction. Total mRNA of femurs and skeletal muscle were extracted. Quantitative real-time PCR was performed to detect mRNA levels of Fzd9 in femur and skeleton muscle.

### Overexpression of Fzd9 in Osteoblasts

A lentivirus vector construction that contains the full-length sequence of rat Fzd9 (GenePharma, Suzhou, China) is used to overexpress Fzd9 in osteoblasts. Briefly, 293T cells were cultured in Dulbecco’s modified Eagles’s medium. The cells were co-transfected with LV5-Fzd9 or LV5-NC (as control) and the packaging vectors (pGag/Pol, pRev, and pVSV-G). The viruses were collected from the culture supernatant 72 h post-transfection, filtered through a 0.45 μm filter, concentrated by centrifugation for 2 h at 20,000 rpm. Titers were determined by infecting 293T cells with serial dilutions of concentrated lentivirus and then counting GFP positive cells after 72-96 h under fluorescent microscopy. After cultured for 2 days, the cells were incubated with lentivirus at 1-1.5×10^6^ TU/ml for 2 days. Polybrene (10 μg/ml) was added to increase the efficiency of lentiviral transduction. The transfected osteoblasts were treated for 3 days with simulated microgravity. The infection rate was evaluated by the expression of green fluorescent protein after incubation with viruses for 2 days. The overexpression of Fzd9 was confirmed by qPCR and Western blotting.

### Two-dimensional clinorotation

A 2D-rotation device (Wheaton, USA) which constantly changes cell orientation by continuous rotation, was used to stimulate certain effects under microgravity as described previously (Yang *et al*, 2013). Briefly, Osteoblasts were seeded at 4×10^4^ cells per flask (12.5 cm^2^, Becton Dickinson; USA). After 24 h, flasks were refilled with DMEM and all air bubbles were carefully removed to prevent fluid shear stress. Flasks for SMG group were fixed on the 2D-rotation device and the rotary speed was 15 rpm/min. The entire device was placed in an incubator in a humidified atmosphere of 5% CO_2_ at 37 °C. Control groups were cultured in the same conditions without clinorotation. Cells were harvested after 1, 2, or 3 days.

### Real-time PCR analysis

Total RNA was extracted using the Trizol Reagent (Invitrogen, USA), and 2μg total RNA was reverse transcribed into cDNA according to the manufacturer’s instructions. Quantitative real-time PCR was performed to detect expression of genes was using the SYBR^®^ Premix Ex Taq kit (Takara, Japan) according to the manufacturer’s instructions and an iQ™5 real-time PCR System (Bio-Rad Laboratories, USA). All reactions were pre-denaturation at 95°C for 3 min, and followed by 40 cycles of 95°C for 10 s, and 60°C for 34 s. After the PCR reaction, melting curve was performed to confirm the specificity and identity of the PCR product. The primers used are presented in Table 1, in which GAPDH was used as endogenous control. Relative gene expression was quantified using the 2^−ΔΔCt^ method.

### siRNA transfection and drug treatment

Small interfering RNAs (siRNAs) targeting YAP was designed and synthesized by GenePharma Corporation (Shanghai, China). Osteoblasts were transfected with Lipo3000 transfection reagent (Thermo Fisher Scientific) according to the manufacturer’s instructions. Osteoblasts transfected with a scrambled non-sense siRNA served as the control (siCon). The siRNA sequences used in the study were as follows: siYAP, 5’-GACATCTTCTGGTCAAAGA-3’ and siCon, 5’-GACATCTTCTGGTCAAAGA-3’. The knockdown efficiency was confirmed by Western blotting.

Osteoblasts were preincubated with the ERK1/2 inhibitor PD98059 (PD, 10 μm, 2 h, Beyotime Biotechnology), Akt inhibitor MK-2206 (2.5 μm, 24 h, Selleck, Houston, TX, USA), actin inhibitor cytochalasin D (cyto D, 1 μm, 2 h, Meilunbio, Dalian, China), actin stabilizer jasplakinolide (Jasp, 50 nM, 2 h, Merck Millipore) or nuclear export inhibitor leptomycin B (LMB, 20 nM, 2 h, Beyotime Biotechnology) and exposed to SMG.

### Alkaline phosphatase (ALP) activity assay and staining

The ALP activity of osteoblasts was analyzed using an ALP assay kit (Nanjing Jiancheng Bioengineering Institute, Nanjing, China) according to the manufacturer ‘s instructions. For ALP staining, osteoblasts were washed with PBS and fixed with 4% paraformaldehydeand then stained using a 5-bromo-4-chloro-3-indolyl phosphate/nitro blue tetrazolium (BCIP/NBT) ALP staining kit (Beyotime Biotechnology, China). The stained cells were then observed and photographed under a microscope and quantified using ImageJ software.

### Western blot

The osteoblasts form each group were harvested and lysed in RIPA buffer (Beyotime Biotechnology, China) containing a protease inhibitor PMSF (Beyotime Biotechnology, China). Equal amounts of protein (30-40 μg) from each sample were subjected to SDS-PAGE and then were transferred to PVDF membranes. The membranes were blocked for 2 h at room temperature using 5% bovine serum albumin (BSA, in Tris-buffered saline with 0.1% Tween-20), and incubated with a primary antibody overnight at 4°C. The antibodies were directed against Fzd9 (1:500, R&D Systems, USA), pGSK3β (1:2000, Cell Signaling Technology, USA), β-catenin (1:2000, BD, USA), pERK (1:500, Santa Cruz, USA), pAkt (1:800, Cell Signaling Technology, USA), GAPDH (1:1000, Good Here, China). The membranes were incubated with secondary HRP-conjugated antibodies for 2 h at RT. Enhanced chemiluminescence (Applygen, Beijing, China) was used to detect the target proteins. ImageJ software was used for densitometric analysis.

### Immunofluorescence

The osteoblasts were gently washed three times with PBS and fixed 20 min at room temperature in 4% paraformaldehyde. The samples were washed 3 times with PBS, permeabilized in 0.2% Triton X-100 for 5-10 min, washed 3 times with PBS, and then blocked with 5% BSA for 2 h at room temperature; incubated with anti-β-catenin (1:200, BD, USA), anti-YAP (1:200, Cell Signaling Technology, USA) and anti-paxillin (1:200, abcam, USA) antibodies at 4 °C overnight; and incubated with secondary antibody conjugated with Alexa Fluor 488-labeled antibodies (1:200, abcam, USA) at 37°C for 1 h. To visualize F-actin, G-actin and cell nuclei, cells were stained with Texas red isothiocyanate-conjugated phalloidin (1:800, Thermo Fischer Scientific, USA), DNase I 488 (1:500, Thermo Fisher Scientific, USA) and 4′,6-diamidino-2 phenylindole (DAPI, Sigma-Aldrich) (1:2000), respectively. Images were captured with confocal microscope (TCS-SP8, Leica Microsystems, Vienna, Austria) or confocal laser-scanning microscope (Andor Dragonfly 500, Leica, England). Images were analyzed by NIH ImageJ software. Z-stack confocal 3D images were obtained with a confocal laser-scanning microscope with a separation interval of 0.15 μm. Z-stack images were analyzed using IMARIS software.

### Transmission electron microscopy (TEM)

For transmission electron microscopy (TEM) experiments, The osteoblasts for SMG group or Con group were fixed with 2.5% glutaraldehyde and 1% paraformaldehyde for 1 h at RT. Cells were post-fixed with 1% osmium tetroxide in 0.8% K4Fe(CN)6 for 1 h at RT in the dark, followed by rinsing with 0.1 M phosphate buffer, dehydration in an acetone series (50%, 70%, 90%, 96%, and 100%; 5 min each), embedment in Epon 812, and polymerization for 48 h at 60 °C. Ultrathin 60-nmthick sections were obtained using a UC7 ultramicrotome (Leica Microsystems). Sections were stained with 5% uranyl acetate for 10 min and lead citrate for 10 min. Sections were collected on 200-mesh grids and examined by TEM (Hitachi, 7065B, Tokyo, Japan) operating at 80 kV, and images were acquired with a charge-coupled device camera (SC1000; Gatan, Pleasanton, CA, USA). Nuclear pores with visible nuclear baskets were selected. The nuclear pore length was measured from one side of the double bilayer, at the point where the nuclear basket starts, to the other side of the double bilayer where the nuclear basket ends.

### Statistical analysis

Data were expressed as the format of means ± standard deviation (s.d.) from at least triplicates. Statistical comparisons were carried out using SPSS 13.0 statistical software with two-tailed Student’s t tests when two cases were compared and with two-way analysis of variance (ANOVA; Tukey or Dunnett) tests when more than two cases were analyzed. Differences were considered to be significant when *p* values were below 0.05. Pearson’s correlation was analyzed, and R^2^ were displayed. Details of sample sizes and significance levels were given in figure legends.

## Data availability

This study includes no data deposited in public repositories.

## Acknowledgements

This work was supported by National Natural Science Foundation of China [Grant Numbers 11972067, 32171310, 11827803, 11421202, U20A20390, 11972068], the 111 Project [Grant Number B13003].

## Author contributions

**Lisha Zheng:** Conceptualization; supervision; funding acquisition; writing-original draft; project administration. **Yubo Fan:** Supervision; funding acquisition; project administration. **Lianwen Sun:** Animal experiments; funding acquisition. **Qiusheng Shi:** Data curation; formal analysis; validation; investigation; visualization; methodology; writing-review and editing. **Jinpeng Gui:** Data curation; formal analysis; investigation; methodology. **Jing Na:** formal analysis; methodology. **Jingyi Zhang:** methodology.

